# An ethologically relevant paradigm to assess visual contrast sensitivity in rodents

**DOI:** 10.1101/2024.03.05.583559

**Authors:** Juan S. Calanni, Marcos L. Aranda, Hernán H. Dieguez, Damian Dorfman, Tiffany M. Schmidt, Ruth E. Rosenstein

## Abstract

In the animal kingdom, threat information is perceived mainly through vision. The subcortical visual pathway plays a critical role in the rapid processing of visual information-induced fear, and triggers a response. Looming-evoked behavior in rodents, mimicking response to aerial predators, allowed identify the neural circuitry underlying instinctive defensive behaviors; however, the influence of disk/background contrast on the looming-induced behavioral response has not been examined, either in rats or mice. We studied the influence of the dark disk/gray background contrast in the type of rat and mouse defensive behavior in the looming arena, and we showed that rat and mouse response as a function of disk/background contrast adjusted to a sigmoid-like relationship. Both sex and age biased the contrast-dependent response, which was dampened in rats submitted to retinal unilateral or bilateral ischemia. Moreover, using genetically manipulated mice, we showed that the three type of photoresponsive retinal cells (i.e., cones, rods, and intrinsically photoresponsive retinal ganglion cells (ipRGCs)), participate in the contrast-dependent response, following this hierarchy: cones ˃> rods ˃>>ipRGCs. The cone and rod involvement was confirmed using a mouse model of unilateral non-exudative age-related macular degeneration, which only damages canonical photoreceptors and significantly decreased the contrast sensitivity in the looming arena.

## INTRODUCTION

Throughout the animal kingdom, the sight of a rapidly approaching object usually signals danger; thus, eliciting a defensive/evasive response. Particularly for rodents, avoiding aerial predators (e.g., hawks, owls) is a central survival function. In that context, Yilmaz and Meister [1] developed a visually-guided behavior test, the “looming test”, for laboratory mice. Looming stimuli are intended to simulate a rapidly approaching aerial predator in the form of a computer-generated, rapidly expanding dark disk on a gray background. In laboratory mice, the looming stimulus triggers a robust running or freezing behavior. This behavioral paradigm was successfully used to identify relevant mouse midbrain visual circuits triggering escape or freeze [2–4]. The superficial superior colliculus (SC), the main retinal synaptic target in rodents, orchestrates the mouse innate defensive responses to visually detected threats [5]. The looming test has been shown to be a robust and reliable vision test across various species, such as mouse [1], [6–7], zebrafish [8], and locust [9]. Although rat models are widely used to recreate specific features of visual diseases, to test therapies at preclinical level, are easy to maintain and testing, and housing is not expensive, scarce information is available about rat behavior in the looming arena. In *Sprague-Dawley* rats, the looming test has been used to assess epilepsy-associated anxiety responses [10], and visual-evoked response in the SC [11], while in *Long-Evans* rats, it has been employed to evaluate the dynamics of hippocampal place cells [12]; however, the looming test has not been yet used as a proxy for rat visual system integrity and function. In this context, we have characterized the influence of sex, age, and daily variation on innate defensive behaviors driven by the looming stimulus. In addition, we tested the effect of a panretinal damage induced by unilateral and bilateral ischemia [13–14] on rat performance in the looming test. Unlike rats, the looming test has been well characterized in mice. The mouse brain retinorecipient areas, and at retina level, OFF-transient αRGCs, W3 RGCs [15–17], and vesicular glutamate transporter 3-expressing amacrine cells [15], [18] have been identified as critical for visually-evoked defensive behaviors; however, there are no data on the influence of disk/background contrast, sex, and age on mouse performance in the looming test. Although it has been shown that retinal photoreceptors participate in the innate fear behavior toward looming stimuli in mice [19], it is still unknown which type/s of photoreceptors is/are involved in the looming stimulus response. Thus, using genetically manipulated mice, we analyzed the involvement of cones, rods, and melanopsin-expressing intrinsically photosensitive RGCs (ipRGCs) in the contrast-dependent looming response. In addition, the performance in the looming arena of mice with non-exudative age-related macular degeneration (NE-AMD), which only affects the outer retina [20–21] was studied.

## MATERIALS AND METHODS

### Animals

All animal procedures were in strict accordance with the ARVO Statement for the Use of Animals in Ophthalmic and Vision Research. The ethic committee of the School of Medicine, University of Buenos Aires (Institutional Committee for the Care and Use of Laboratory Animals, (CICUAL)) and the Animal Care and Use Committee at Northwestern University approved this study, and all efforts were made to minimize animal suffering. Male and female *Wistar* rats, and male and female C57BL/6J mice were bred in house and kept under controlled temperature, luminosity, and humidity, and under a 12-h light/12-h dark lighting schedule. Young male rats and mice (2-3 months old) were used as controls, whereas 13-14 months old rats and mice were used as aged animals. Experimental sessions were carried out between ZT 4.00 and ZT 6.00 at day-time, while night time experiments were performed between ZT 14.00 and ZT 16.00. To ascertain the specific contribution of individual photoreceptors in the looming response, transgenic mice originally employed by Altimus et al. [22] to selectively eliminate rods, cones and/or ipRGCs, without the induction of non-specific retinal degeneration were utilized. The study encompassed sex- and aged-matched genotypes, as follows: C57Bl/6J wild type (Opn4^+/+^, Gnat1^+/+^, Gnat2^+/+^, Control), rod knockout (Opn4^+/+^, Gnat1^-/-^, Gnat2^+/+^, RKO), rod and melanopsin knockout (Opn4^-/-^, Gnat1^-/-^, Gnat2^+/+^, or cone-only, (C only)), cone knockout (Opn4^+/+^, Gnat1^+/+^, Gnat2^-/-^, CKO), cone and melanopsin knockout (Opn4^-/-^, Gnat1^+/+^, Gnat2^-/-^, or rod only, (R only)), cone and rod knockout (Opn4^+/+^, Gnat1^-/-^, Gnat2^-/-^, or melanopsin only, (M only)) and rod, cone, and melanopsin knockout (Opn4^-/-^, Gnat1^-/-^, Gnat2^-/-^, or triple knockout, TKO) mice.

### Looming arena

The experimental arena consisted of a Plexiglas box with a monitor embedded in the ceiling. The arena dimensions were 70×70×70 cm^3^ for rats, and 40×40×40 cm^3^ for mice. Throughout the experimental sessions, the monitor displayed a dim gray background, on which a dark expanding disk was presented as the looming stimulus (Figure 1A). Experiments were performed in a mesopic environment.

**Figure 1.**
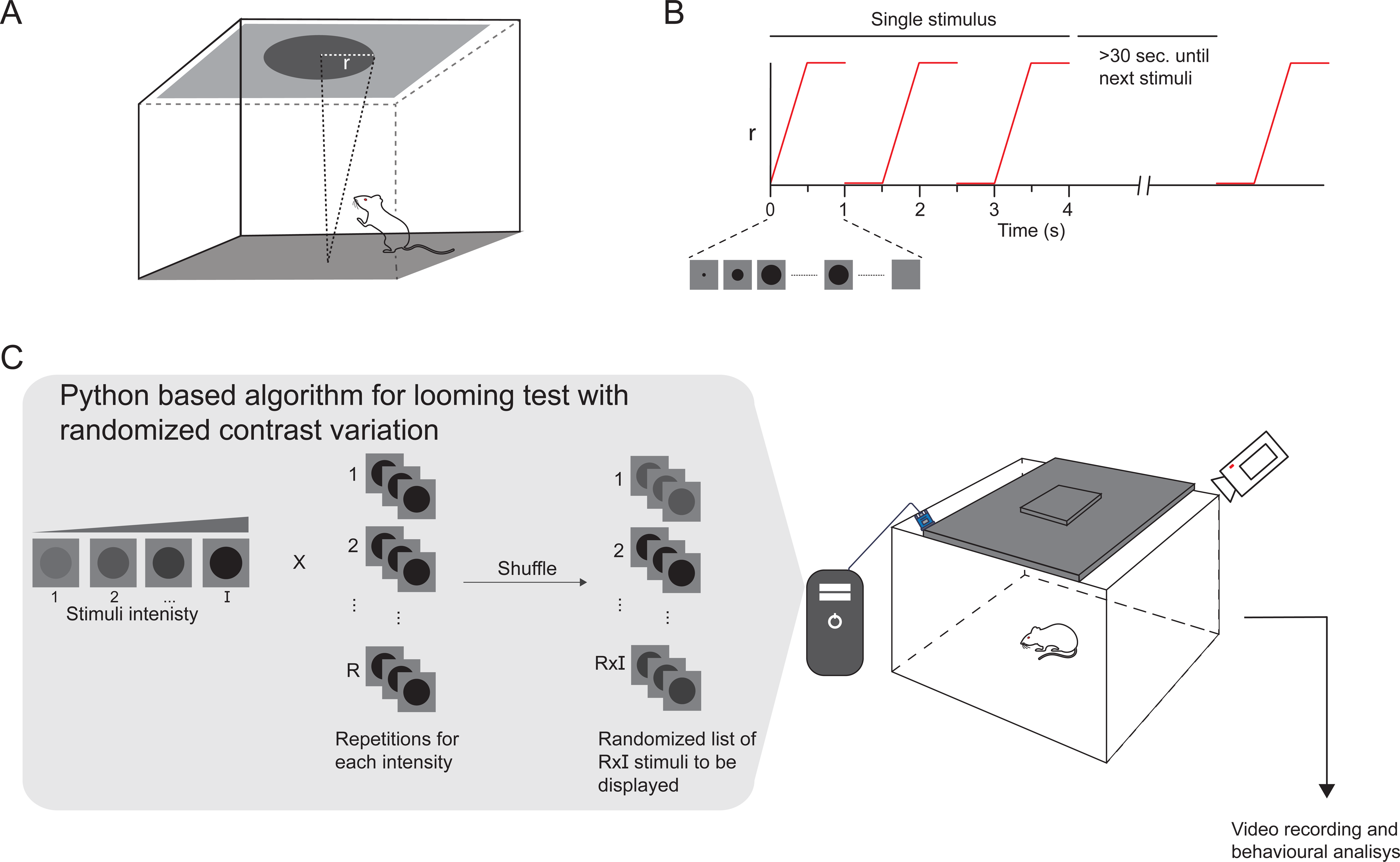
Protocol for studying the influence of disk/background contrast on the looming response. Schematic of the looming arena with the monitor embedded in the ceiling and displaying one disk expansion. B) Schematic representation of the looming stimuli progression. C) Algorithm for looming test with randomized contrast variation, which consisted in a publicly available python-based software designed to display RxI looming stimuli in a randomized way, where I represents the number of disk/background contrast magnitude to be displayed, and R represents the repetitions of each contrast magnitude. The values for R and I were defined by the operator, prior to initiating the test.

### Stimulus dynamics

Each stimulus consisted of three expansions of a dark disk projected onto the gray background of a PC monitor. A single disc expansion consisted of two phases: 1) the disk expanded from r = 0 to r max over a duration of 0.5 seconds, 2) then, the disk remained static at its maximum expansion (r max) for 0.5 seconds before disappearing. A 0.5-second interval was left before the next disk expansion; thus, each stimulus presentation lasted 4 seconds. At least 30 seconds were left between consecutive stimulus presentations (Figure 1B).

### Disk/background contrast variations

A publicly available Python-based algorithm (https://github.com/SalvadorCalanni/LTCV) was used to present dark expanding disks with different contrast levels in a random fashion. The contrast level was assessed by measuring the irradiance, using a photometer positioned at the center of the arena. Specifically, the irradiance of the fully expanded grey disk (Id) was compared with the irradiance of the background alone (Ib). The Michelson index (MI) was then calculated to quantify the contrast level as follows:

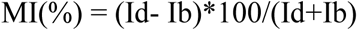

A range of contrasts was constructed between the lowest contrast that did not provoke response in any control animal, and the highest contrast that, consistently, elicited response in all control animals. The number of contrasts (C), and the repetitions of each one of them (R) were predetermined before the looming sessions, which resulted in a RxC list of stimuli that were randomly presented to animals during the experimental session. The order of contrast presentations was randomized to minimize any potential order effects, and animal habituation (Figure 1C). The RxC number of looming stimuli was determined based on the duration of animal exploratory behavior. Exploratory activity lasted approximately 30 minutes for rats, and around 45 minutes for mice. The highest number of contrasts and repetitions were included within these periods; specifically, rats were exposed to 5 contrast levels, each repeated 3 times (i.e., a total of 15 stimuli), while mice were exposed to 6 different contrast intensities, each one repeated 4 times (i.e., a total of 24 stimuli), resulting in experimental sessions that lasted approximately 20 minutes for rats and 35 minutes for mice. Each experimental session was recorded with a video camera for later analysis.

### Experimental sessions

Animals were habituated to the experimental room for 2 h. No food or water restrictions were imposed before the tests. Animals were gently placed in the arena, and allowed to acclimate for 5 minutes before the first stimulus presentation. Each stimulus was manually triggered by an operator, only when the animal stanned close to the arena center.

### Behavioral analysis

Video recordings of experimental sessions were analyzed by three independent and masked observers. The analysis focused on identifying stereotyped behaviors in response to looming stimuli.

### Retinal ischemia in rats

Young male *Wistar* rats were anesthetized with ketamine hydrochloride (150 mg/kg) and xylazine hydrochloride (2 mg/kg) administered intraperitoneally. After topical instillation of proparacaine, the eye anterior chamber was cannulated with a 30-gauge needle connected to a pressurized bottle filled with sterile normal saline solution. Retinal ischemia was induced by increasing intraocular pressure to 120 mmHg for exactly 40 min in one eye, as previously described [13–14]. Complete ocular ischemia, characterized by flow cessation in retinal vessels as shown by funduscopic examination, was achieved. During and after this manipulation, animals were kept normothermic with heated blankets. The contralateral eye remained intact, because in previous works we showed that the sham procedure (i.e., eyes cannulation without raising IOP), did not affect retinal function and histology [13–14]. A group of animals were submitted to bilateral ischemia. A few animals in which cataracts developed due to lens injury, were not used any further in the experiments.

### Experimental non exudative age-related macular degeneration in mice

Unilateral NE-AMD was induced through ipsilateral superior cervical ganglionectomy (SCGx), as previously described [20–21]. Briefly, the left superior cervical ganglion (SCG) was surgically removed under aseptic conditions, resulting in the ipsilateral loss of sympathetic innervation, while a unilateral sham procedure was performed in control animals. The neck incision was closed with nylon sutures and mice recovered without complications. This model mimics many of NE-AMD features in humans, such as choroidal thickening, Bruch’s membrane thickening, photoreceptor and retinal pigment epithelium cell dysfunction and death, without affecting the inner retina [20-[21].

### Electroretinography

Electroretinographic activity was assessed at 4 weeks after ischemia in rats and 10 weeks after SCGx in mice, as previously described [13–14],[20–21]. Briefly, after 6 h of dark adaptation, animals were anesthetized under dim red light. Phenylephrine hydrochloride and tropicamide were used to dilate the pupils, and the cornea was intermittently irrigated with balanced salt solution to maintain the baseline recording and to prevent keratopathy. Animals were placed facing the stimulus at a distance of 20 cm. All recordings were completed within 20 min and animals were kept warm during and after the procedure. A reference electrode was placed through the ear, a grounding electrode was attached to the tail, and a gold electrode was placed in contact with the central cornea. A 15W red light was used to enable accurate electrode placement, without affecting dark adaptation and was switched off during the electrophysiological recordings. Electroretinogram (ERG) recordings were made with a HMsERG model 2000 (Ocuscience LLC), equipped with a Ganzfield dome fitted with a white light-emitting diode stimulus at a distance of 2 cm from the eye. To assess scotopic ERG a-wave and b-wave, 15 full-field flashes separated by a 10 s interval (flash intensity 10 cd.s.m^−2^) were averaged.

### Histological Evaluation

Four weeks after ischemia or six weeks after SCGx, animals were anesthetized and intracardially perfused with saline solution, followed by a fixative solution containing 4% formaldehyde in 0.1 mol/L PBS (pH 7.4). Then, the eyeballs were carefully removed and immersed for 24 hours in the same fixative. After dehydration, eyes were embedded in paraffin wax and sectioned (5 µm) along the vertical meridian through the optic nerve head. Rat retinal sections from ischemic and control eyes were stained with hematoxylin and eosin. Mouse retinal sections from SCGx and control eyes were stained with Liliés trichrome. Microscopic images were digitally captured with a microscope (Eclipse E400, Nikon, Tokyo, Japan); 6-V halogen lamp, 20 W, equipped with a stabilized light source) and a camera (Coolpix s10; Nikon; Abingdon, VA, USA) and analyzed by masked observers.

### Statistical analysis

To evaluate the relationship between stimulus contrast and response, the proportion of positive responses (including all stereotyped responses) was measured for each contrast. Subsequently, the influence of contrast magnitude, age, sex, time of the day, rat unilateral or bilateral ischemia, mouse genotypes, and mouse with experimental NE-AMD, on the probability of eliciting a defensive response was assessed using a binomial generalized linear model (GLM). The GLM analysis was conducted in R statistical software, treating contrast level, age, sex, time of the day, rat unilateral ischemia or bilateral ischemia, mouse genotype, and mouse NE-AMD as fixed factors. The disk/background contrast necessary to elicit a 50% positive response (C50) for each animal was predicted using the GLM. Differences in C50 between groups was analyzed using the t-student’s test after confirming homoscedasticity assumption through Bartlett’s test. All data and statistical analysis were upload to https://github.com/SalvadorCalanni/LTCV.

## RESULTS

Figure 2A shows behavioral responses of visually intact young adult male *Wistar* rats in the looming arena, under different disk/background contrasts. Using identical duration, stimulus size of the dark disk, and expansion dynamics, three distinct behavioral responses were observed in rats: 1) “head bobbing”, which involved lateral movements of the head, 2) “upward rearing” where the rat stood on its hind legs while sniffing/observing the ceiling, and 3) “freezing”, characterized by the sudden cessation of arena exploration, remaining motionless for a few seconds. Since fleeing to a shelter was not observed in any rat, the shelter was removed from the arena. To better understand contrast influence on looming response, the proportion of responses for each contrast was calculated. At high disk/background contrasts, freezing and upward rearing were more frequent, while at low/middle contrasts, head bobbing prevailed (Figure 2A). Given that response values could range from 0 to 1, we employed a GLM of the binomial family to model the frequency of responses as a function of contrast magnitude. Our analysis revealed that contrast magnitude significantly explained the variation in responses, indicating a well-fitted sigmoid relationship (P<0.0001), as shown in Figure 2B. The influence of sex, age, and time of the day on contrast magnitude-dependent response is shown in Figure 3. To enhance the representation of intra-group variations, the GLM was employed to compute the C50 for each individual animal, as depicted in Supplementary Figure 1. For young adult female *Wistar* rats, data also adjusted to a sigmoid contrast-response relationship, but the contrast needed to elicit a response was lower for females according to the GLM predictions, as indicated by the leftward shift in the GLM predictions curve (P<0.05). This is further supported by the significantly higher C50 observed in young adult males compared to young adult females (Figure 3A). Moreover, the GLM predictions curve shifted to the right for old males compared to young males (P<0.01), and the C50 for old males was significantly higher than that for young adult males (Figure 3B). In young adult male rats registered at ZT 14.00-16.00, data also adjusted to a sigmoid relationship between the contrast magnitude and response with the curve shifted to the left compared to males registered at noon (P<0.01). while the C50 was significantly lower than that observed at ZT 4.00-6.00 (Figure 3C). In all these experimental groups, a 100% of response was reached with the highest contrast examined.

**Figure 2.**
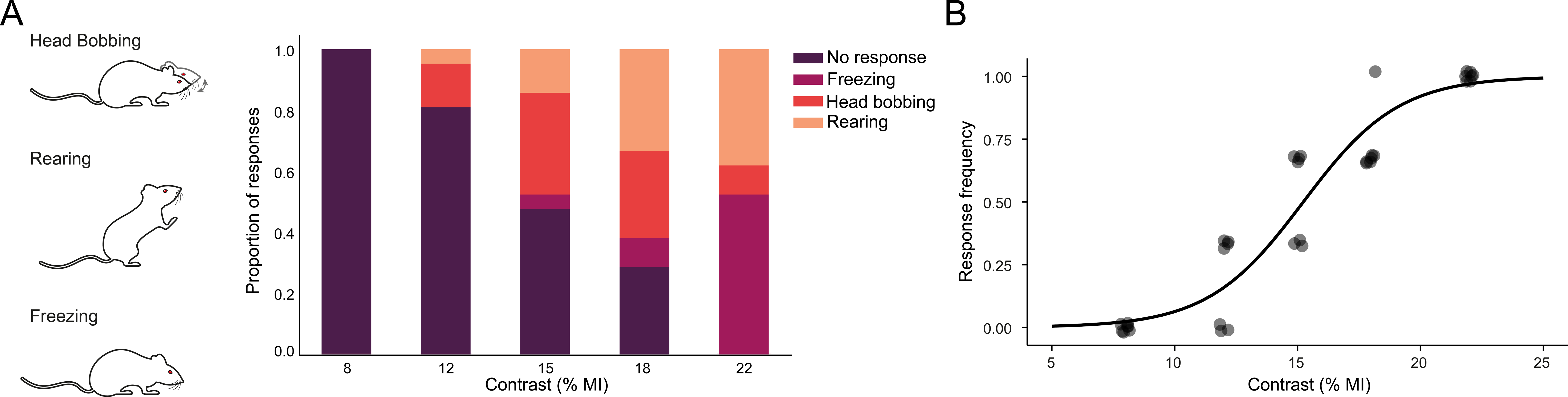
Rat response to different disk/background contrasts. A) Schematic representation of different stereotyped responses in rats, along with the distribution of the responses versus contrast magnitude for young male rats. B) Response frequency as function of the contrast magnitude for young male rats. The curve depicts predictions calculated by the GLM.

**Figure 3.**
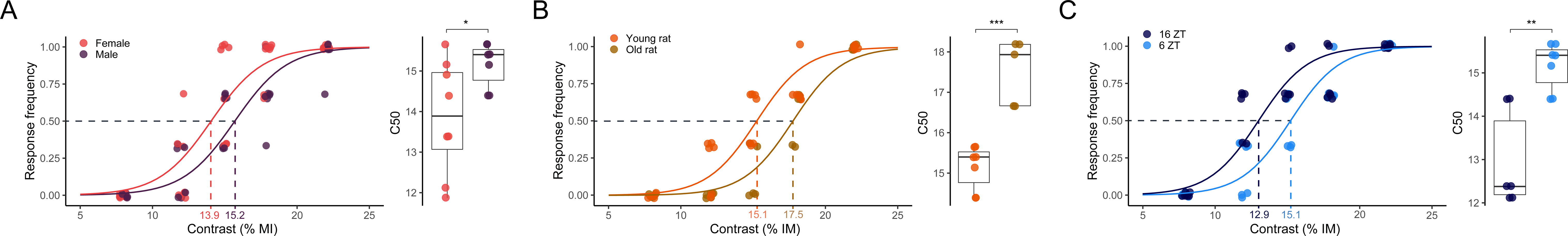
Influence of sex, age, and time of the day on rat contrast-dependent response. Response frequency as function of the contrast magnitude and individual C50 value in: A) young male **and young female rats, B) young** male **and old** male **rats, C) young** male **rats at ZT 4.00 - 6.00 and ZT 14.00-16.00. In the response frequency vs. contrast plots, each point represents the proportion of responses of an individual animal for a specific contrast magnitude, and the line represents the GLM predictions for the entire group. Group C50 is also indicated in the plot. In the C50 plots, each point represents the C50 of one single animal. *P< 0.05, **P< 0.01, ***P< 0.001**.

In order to know whether a panretinal damage affects rat behavior in the looming arena, young adult male rats were submitted to unilateral or bilateral ischemia. Unilateral or bilateral ischemia induced a significant decrease in flash scotopic ERG a- and b-wave amplitude, and notorious structural alterations with a decrease in retinal layer thickness at 4 weeks post-ischemia (Figures 4B, 4C). No differences were observed between unilateral and bilateral ischemic damage on the retinal function and structure (data not shown). Notwithstanding, a sigmoid relationship between the contrast magnitude and response was also observed in rats submitted to unilateral or bilateral ischemia, but with the GLM prediction, curves shifted to the right (P<0.0001), and the resulting C50 were: bilateral ischemia > unilateral ischemia > intact animals (Figure 4D).

**Figure 4.**
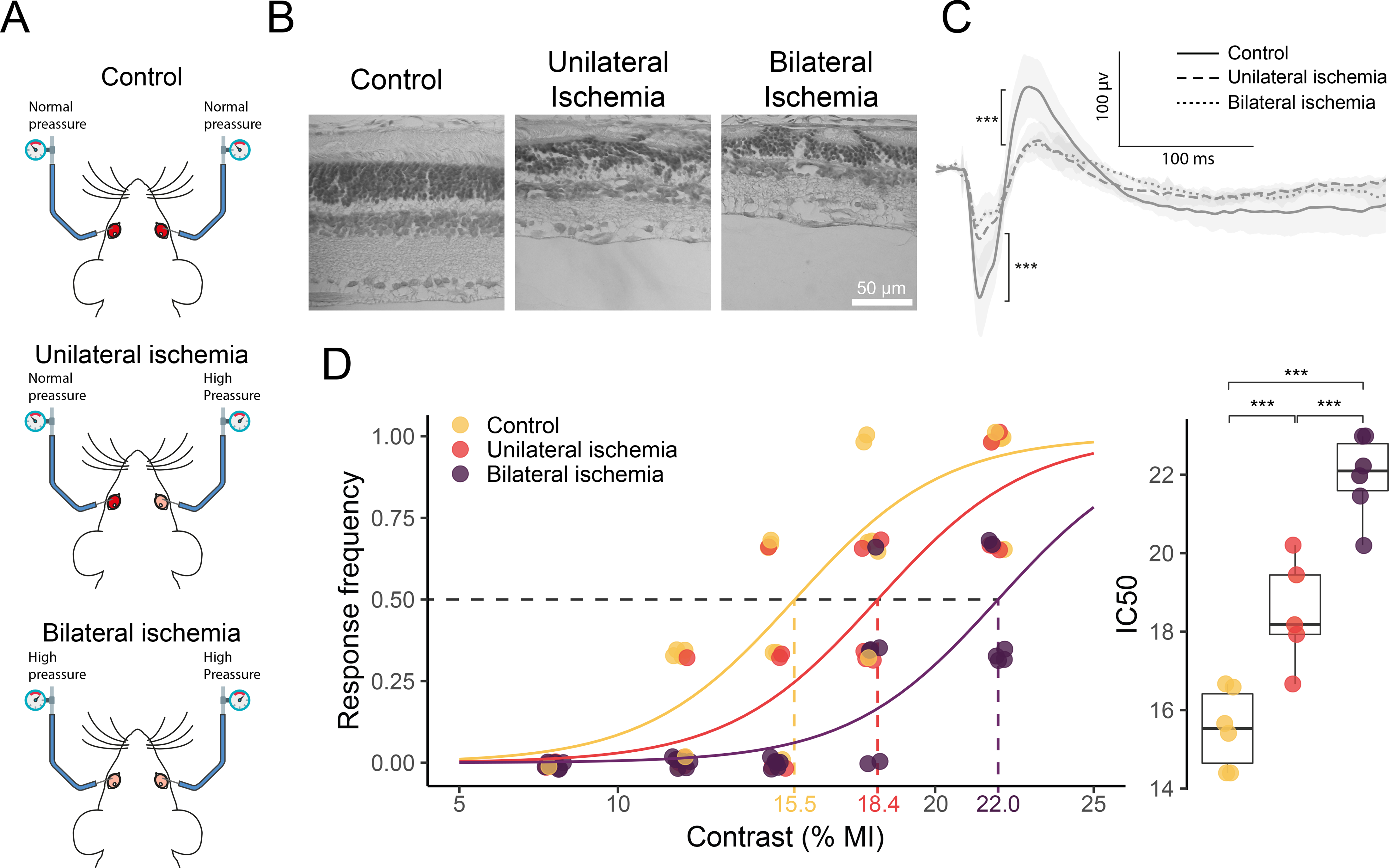
Effect of unilateral or bilateral ischemia on rat contrast sensitivity. A) Schematic representation of the surgical procedure for sham treatment or ischemia. B) Representative photomicrographs (H&E); C) Recordings of scotopic ERGs from sham-treated, or rat submitted to unilateral or bilateral ischemia. D) Response frequency as function of the contrast magnitude and individual C50. In the response frequency vs. contrast plots, a point represents the proportion of responses of an individual animal for a specific contrast magnitude, and the line represents the GLM predictions for the entire group. The group C50 is also indicated in the plot. In the C50 plots, each point represents the C50 of one single animal. ***P< 0.001.

Although innate fear response evoked by looming stimulus have been extensively studied in mice, the influence of sex and age in C57BL/6J mice has not been examined. Consistently with rat responses, mice exhibited upward rearing and freezing (but not head bobbing) in response to different contrasts. Moreover, mice displayed an additional response: running, (i.e., a rapid sprint towards one of the arena corners) immediately after the looming stimulus (Figure 5). The relative frequency of these stereotyped responses depended on the contrast magnitude, as shown in Figure 5A. At high contrasts, the most frequent response was freezing, while rearing prevailed at low contrasts. Computing responses as previously described for rats, behavioral data adjusted to a sigmoid-type contrast-response relationship also in wild-type mice (P<0.0001) (Figure 5B). The contrast-dependent mouse performance in the looming arena was also influenced by sex and age, as shown in Figure 5B. The C50 was significantly higher for young adult males than for young adult females, for aged males than for young adult males, and for aged females than for young adult females, whereas it did not differ between old males and old females. In order to identify retinal photoreceptor type/s involved in contrast-dependent looming performance, knockout mice were used. For this purpose, sex-and aged-matched, control, RKO, C only, CKO, R only, M only, and TKO genotypes were tested under different contrast magnitudes, as shown in Figure 6. The contrast-dependent response in genotypes preserving intact cones (i.e., RKO and C only) did not differ from control mice, whereas animals lacking cones showed an increase in C50. In particular, CKO and R only mice needed a higher contrast magnitude to reach a response frequency similar to that from animals with intact cones. M animals, lacking both cones and rods, incipiently responded at highest contrasts tested, whereas animals lacking any type of retinal photoreceptors did not respond at all. The influence of retinal damage induced by unilateral NE-AMD in wild-type mice on the performance in the looming arena is shown in Figure 7. Experimental NE-AMD only affected photoreceptor function, as shown by a significant decrease in the ERG a-wave amplitude, without changing ERG b-wave amplitude. At structural level, photoreceptor outer segment loss and outer nuclear layer disorganization were evident (Figure 7A). Unilateral NE-AMD significantly increased the C50 (Figure 7B) as compared with young adult male mice submitted to a unilateral sham procedure.

**Figure 5.**
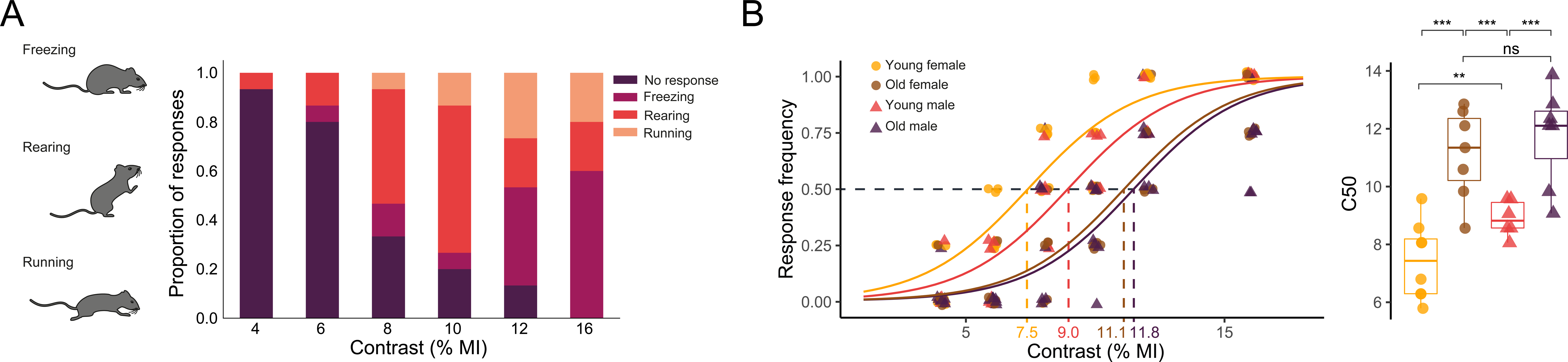
Mouse responses to different disk/background contrasts. A) Schematic illustration of the different stereotyped responses in mice, along with the distribution of the responses versus contrast magnitude for young male mice. B) Response frequency as function of the contrast magnitude and individual C50 for young male, young female, old male and old female mice. In the response frequency vs. contrast plots, a point represents the proportion of responses of an individual animal for a specific contrast intensity, and the line represents the GLM predictions for the entire group. The group C50 is also indicated in the plot. In the C50 plots, each point represents the C50 of one single animal. **P< 0.01, ***P< 0.001.

**Figure 6.**
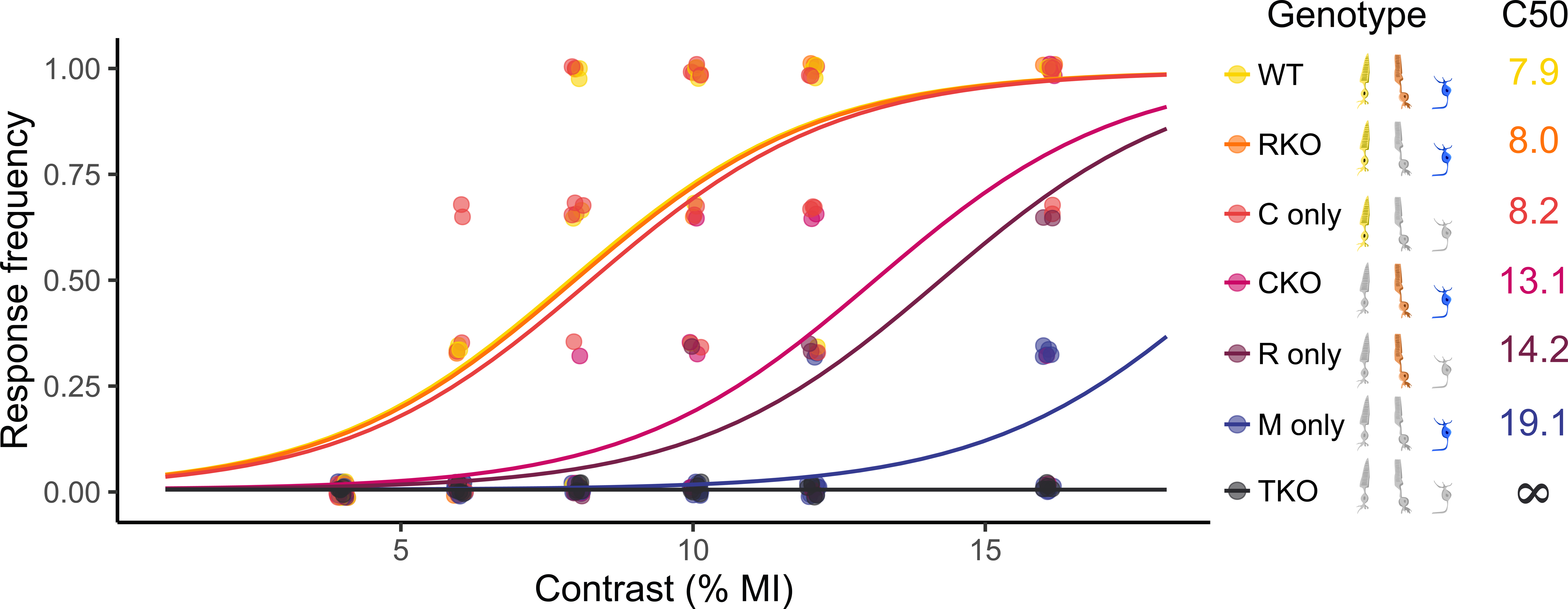
Analysis of different photoreceptor type involvement in the contrast-dependent response. Response frequency as function of the contrast magnitude for mice with different genotypes are shown. Each point represents the proportion of responses of an individual animal for a specific contrast magnitude, and the line represents the GLM predictions for the entire group. The group C50 is numerically indicated with the genotype reference.

**7.**
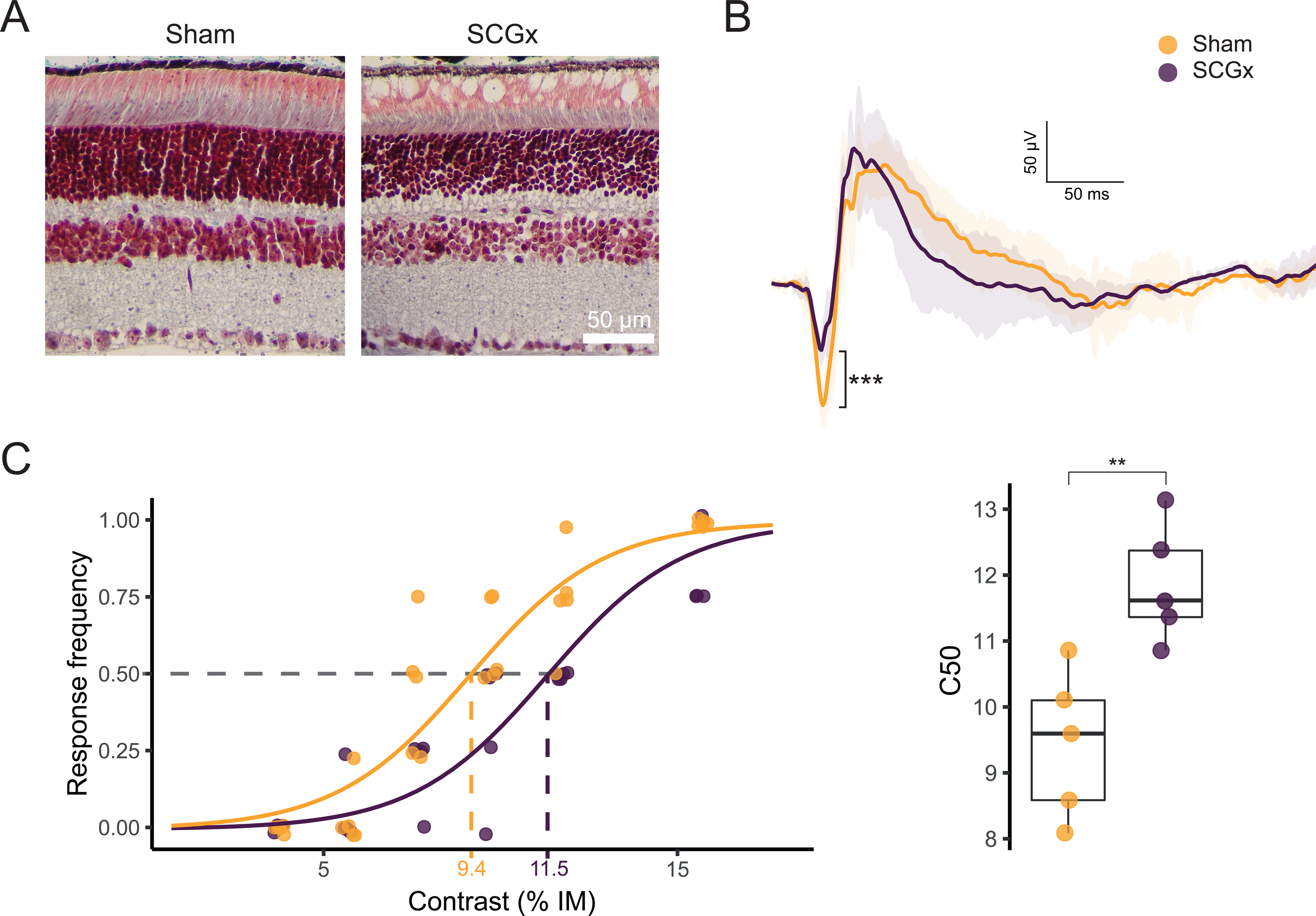
Effect of experimental NE-AMD induced by unilateral SCGx on contrast-dependent response in the looming arena. A) Scotopic ERG amplitude for control and SCGx mice. SCGx significantly decreased ERG a-wave amplitude. B) Response frequency as function of the contrast magnitude and individual C50. In the response frequency vs. contrast plots, a point represents the proportion of responses of an individual animal for a specific contrast intensity, and the line represents the GLM predictions for the entire group. The group C50 is also indicated in the plot. In the C50 plots, each point represents the C50 of one single animal. **P< 0.01, ***P< 0.001.

## DISCUSSION

The laboratory rat has become one of the major animal models in preclinical ophthalmologic studies. In that context, although visually-guided behavior tests are strong tools to evaluate rodent vision, the looming test has not been previously used in rats as a visual function index. Young adult male *Wistar* rats challenged with overhead looming stimuli showed different defensive response types. The contrast information of the looming stimulus biases the type of defensive behavior in goldfish (*Carassius auratus*) [23], but to our knowledge, the effect of changing the dark disk/grey background contrast in the looming test has not been previously analyzed in rodents. As shown herein, the relative contribution of rat behavioral response types depended on the disk/background contrast magnitude, which could suggest that specialized channels are involved in facing different threatening visual scenarios. Rat head bobbing has been described in different experimental conditions [24–25], and has been associated to hyperkinesis [26]. In the looming arena, rat head bobbing occurred as an innate defensive behavior mostly at low/middle contrasts, and probably as a strategy to increase the visual depth perception against an aerial threat. To focus on retinal processing, and to avoid inter- and intra-individual variability in innate behaviors, the response was assessed in an all-or-none fashion, as an indicator of whether a rat detected or not a particular disk/background contrast. In addition, rats were presented with multiple repeats of the same or different disk/background contrasts selected at random, to avoid a possible bias due to habituation [3]. The global analysis including any type of defensive response of young adult male *Wistar* rats against a range of disk/background contrasts adjusted to a saturation sigmoid function that reached a plateau in which 100% of animals responded positively. An adjustment to a sigmoid-like contrast-response curve was also observed in old males, and young adult females, but the curve shifted to the right in aged male rats, and to the left in young adult females, as compared to young adult males, supporting a higher contrast sensitivity in young adult female rats. In agreement, a greater innate fear in response to looming stimuli has been described in female *Sprague Dawley* rats [27]. Being nocturnal animals, rats are more active at the dark phase than during the light phase, which could account for a more sensitive response at the active than at the rest phase, as shown by a higher C50 at noon than at midnight. In the same line, a poorer vision in old male rats [28] correlated with a higher C50 than that in young adult rats. At present, we cannot formally discern the relative contribution of general activity and age-dependent influence on the contrast sensitivity, but with strain-, age- and sex-matched groups, and testing animals at the same time of the day, the C50 could be used as a proxy for contrast sensitivity. Acute retinal ischemia provokes a significant rod and cone dysfunction (shown by flash electroretinography) [13–14], and increased the C50 as compared with visually intact young adult rats, which could suggest the involvement of classical photoreceptors in the looming response. However, since retinal ischemia also decreases canonical RGC number, without affecting ipRGCs number and function [29], and induces a deficit in the retina-SC communication [13–14], the involvement of canonical RGCs-SC circuit alteration on the retinal ischemia-induced changes in the C50 cannot be ruled out. Although unilateral and bilateral ischemia induced a similar retinal damage, the C50 was significantly higher in rats with bilateral than with unilateral ischemia, supporting that animals with only one intact retina were able to detect the looming stimulus, although with less sensitivity than those with both intact retinas.

Although the influence of the angular size and the expanding speed of the looming stimulus on the evasive behavior has been studied [1], [7], [30], no information on the disk/background contrast-dependent response in mice has been provided. As with rats, mice showed a mixed defensive behavior that varied in a contrast-dependent manner. At middle contrasts, the most frequent response was upward rearing, while at high contrasts, freezing and running predominated, consistently with the perception of a higher proximity of an aerial menace. Behavioral data also adjusted to a sigmoid contrast-response relationship in adult male wild type mice, which was influenced by sex and age, in a similar way to that observed in rats (i.e., old males = old females > young males > young females). Systemic application of N-methyl-N-nitrosourea that removes retinal photoreceptors, provokes the disappearance of the innate fear behavior toward looming stimuli in mice [19], but there is still no information on which type/s of retinal photoreceptors are involved in the contrast-dependent looming response. To gain insight into this aspect, genetic manipulations in mice (more amenable to genetic perturbations than rats) were used. The response of mice preserving cones (RKO and C only) did not differ from control mice, whereas animals lacking cones were much less responsive. M only animals incipiently responded at highest contrasts, and in animals lacking cones, rods, and mRGCs, none response was evident at any contrast.

Taken together, the present results support that in different ways, retinal cells with intrinsic photoreceptive capacity participate in the contrast-dependent looming response, following this hierarchy: cones ˃> rods ˃>>ipRGCs. In fact, in an experimental model of NE-AMD in mice, which unlike the panretinal ischemic damage, only affects the outer retina (i.e., photoreceptors and retinal pigment epithelium), with a complete preservation of the inner retina [20–21], the C50 significantly increased as compared with sham-treated mice. At first glance, it could seem surprising that nocturnal animals with rod-dominant retinas, foraging for food and facing predators at the sunset/moonlight night, mostly depended on cones for a defensive behavior against an aerial prey. Each lifestyle and habitat have contributed to evolutionarily select the diversity of photoreceptor arrangements in mammals [31]. In the mouse, there are two spectral cone types: one expressing an opsin sensitive to short-wavelengths (S-opsin), and the other, to middle-to-long-wavelengths (L-opsin), although most of cones co-express both opsins [32–34]. Dual cones broaden the spectral range, allowing a better vision in varied spectral compositions of ambient light [31–32]. The S-opsin in the ventral retina encodes preferentially dark contrast; thus, cones expressing S-opsin whose highest densities are located in the ventral retina could be sky sensors for aerial preys, while the L-opsin in the dorsal retina is used to see the ground [35]. In addition, since center/surround antagonism requires the cone circuitry, cones involvement in behavior at mesopic intensities seems plausible. Moreover, even in the natural environment, the light intensities in moonlight and at dusk would not be below the mesopic range, where both rods and cones can contribute. In fact, although with less sensitivity than C only and RKO mice only, rods (R only) also contributed to innate defensive response; therefore, alterations affecting rods may also being detectable when studying the relationship behaviors vs. contrasts. The rod signal is fed into the cone system after a detour, involving the rod bipolar and AII amacrine cells, producing appropriate signal polarities for the ON and OFF pathways. Rod signals also enter the cone system through two other pathways; rods can drive neighboring cones directly through electrical junctions, making connections with an OFF bipolar cell that services primarily cones. Once the rod signal has reached the cone bipolar cells through these pathways, it can take advantage of the same intricate circuitry of the inner retina. Therefore, the lower response in rod-retinas could be attributed to a lower rod contrast sensitivity or to an obliteration of rod-cone dialogue. Since M only animals showed a defensive behavior only with the highest contrast tested, it seems that ipRGCs could trigger innate defensive responses against high disk/background contrasts.

In the present report, using the innate response to the looming stimulus, we developed a new test to measure contrast sensitivity both in rats and mice by varying disk/background contrast (i.e., the looming test with contrast variation (LTCV)). In addition, we have gained insight in the relative contribution of retinal photoreceptive cells to the contrast-dependent response in the LCTV. Using the LCTV, animals can be individually evaluated in a relatively short term, and provides a quantitative measurement (i.e., the C50) that can be used as an index of contrast-sensitivity in rodents. In comparison with other methods for the assessment of visual function intactness, several advantages support the suitability of the LTCV, as follows: 1) unlike the ERG, it does not requires anesthesia; 2) unlike the optokinetic reflex, it does not need surgery to attach a fixation device to the skull [35]; 3) unlike the optomotor response, it does not requires the experimenter training to subjectively detect subtle mouse head movements; 4) unlike the visual water task, rodents do not require a pre-training phase of learning; 5) it has low cost and requires a very simple setup, with which visual stimuli are simple to generate, precise, and highly controllable; 6) is able to reveal an altered contrast-dependent response even when only one eye was affected. Therefore, the LTCV could be a non-invasive tool to test new experimental models of visual impairment in rodents, as well as to evaluate the efficacy of therapeutic treatments. Future experiments should include different rat and mouse strains, and analyze whether this test can be used to discriminate levels of vision loss in other experimental models of retinal diseases.

## Acknowledgments

This research was supported by grants from the Agencia Nacional de Promoción Científica y Tecnológica [PICT 0415, PICT 0157, PICT 1506]; The University of Buenos Aires [20020220100070BA]; Consejo Nacional de Investigaciones Científicas y Técnicas [PIP 12320220100606CO], Argentina. DP2 EY022584 and R01 EY030565 awarded to T.M.S by the National Institute of Health, United States. The funding organizations have no role in the design or conduct of this research. The authors report no conflicts of interest.

## • Author contributions

J.S.C. designed, performed, and analyzed all experiments H.H.D developed experimental models, performed immunohistochemical analysis, electroretinography D.D. revised critically the manuscript M.L.A; T.M.S., and R.E.R. coordinated the project, analyzed the data, and wrote the manuscript.

## • Data availability statement

Data available at the following link: https://github.com/SalvadorCalanni/LTCV.

---

**Supplementary Figure 1. Example of the C50 calculation for an individual animal.** Overall and individual predictions for 4 different rats are depicted in the response frequency vs. contrast plot. The individual curves were utilized to calculate the C50, as illustrated in the Figure. These values were subsequently plotted in the C50 plot on the right.

