## Supplementary figures and images for "An ethologically relevant paradigm to assess visual contrast sensitivity in rodents"

### Supplemental Figure 1

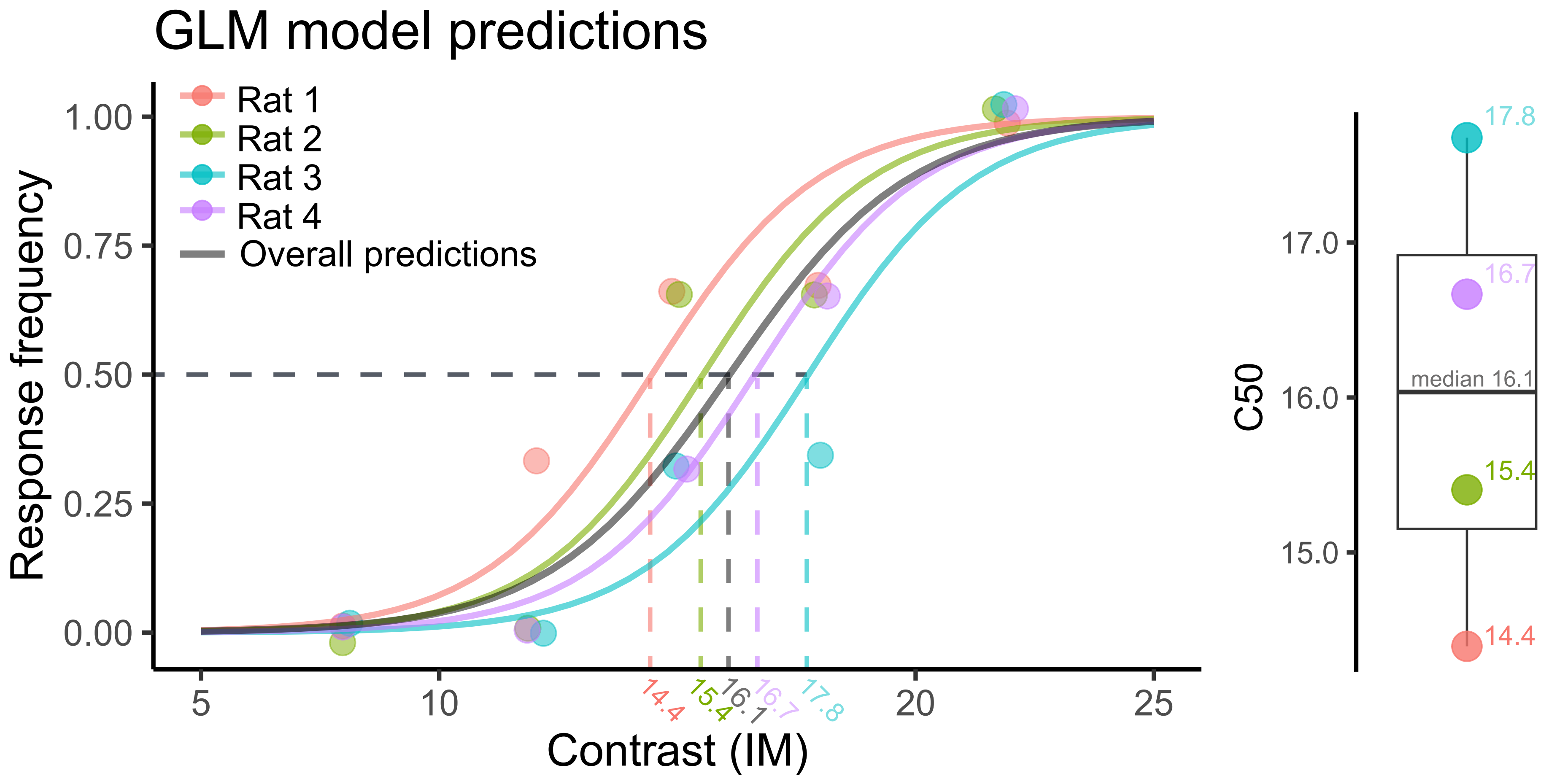
